# Effect of human synovial fluid from osteoarthritis patients and healthy individuals on lymphatic contractility

**DOI:** 10.1101/2020.12.02.408294

**Authors:** Eleftheria Michalaki, Zhanna Nepiyushchikh, Fabrice C. Bernard, Josephine M. Rudd, Anish Mukherjee, Jay M. McKinney, Thanh N. Doan, Nick J. Willett, J. Brandon Dixon

**Author notes:** These authors contributed equally to this work (Eleftheria Michalaki, Zhanna Nepiyushchikh). Correspondence and requests for materials should be addressed to N.J.W. and J.B.D.

## Abstract

The lymphatic system has been proposed to play a crucial role in the development and progression of osteoarthritis (OA). The synovial fluid (SF) of arthritic joints contains mediators of the inflammatory response and products of the injury to articular tissues, while lymphatic system plays a critical role in resolving inflammation and overall joint homeostasis. Despite the importance of both the lymphatic system and SF in OA disease, their relationship is still poorly understood. Here, we utilized SF derived from osteoarthritis patients (OASF) and healthy individuals (HSF) to investigate potential effects of SF on migration of lymphatic endothelial cells (LECs) *in vitro*, and lymphatic contractility of femoral lymphatic vessels (LVs) *ex vivo*. Both OASF and HSF treatments led to an increased migratory response *in vitro* compared to LECs treatment with media without serum. *Ex vivo*, both OASF and HSF treatments to the lumen of isolated LVs led to significant differences in the tonic and phasic contractions and these observations were dependent on the SF treatment time. Specifically, OASF treatment transiently enhanced the RFLVs tonic contractions. Regarding the phasic contractions, OASF generated either an abrupt reduction after 1 hr of treatment or a complete cease of contractions after an overnight treatment, while HSF treatment displayed a gradual decrease in lymphatic contractility. The observed variations after SF treatments suggest that the pump function of lymphatic vessel draining the joint could be directly compromised in OA and thus might present a new therapeutic target.

## Introduction

Osteoarthritis (OA) is the most common form of arthritis affecting millions of people worldwide, with notably more than 3 million cases per year in the US (Arden and Nevitt, 2006; Chen *et al.*, 2016; Cucchiarini *et al.*, 2016). OA can damage any joint but hands, knees, hips, feet, and spine are the most commonly affected areas (Iolascon *et al.*, 2017). Symptoms of OA include pain, joint dysfunction, and deformity, while gender, age, obesity, and the presence of biomechanical predisposing factors seem to directly affect the development of OA (Cucchiarini *et al.*, 2016; Maricar *et al.*, 2016; Berenbaum *et al.*, 2018; Bortoluzzi, Furini and Scirè, 2018; Shu *et al.*, 2020). OA is characterized by the gradual degradation of the articular cartilage, along with changes in the subchondral bone, synovium, meniscus, tendons, and muscles (Robinson *et al.*, 2016; Mathiessen and Conaghan, 2017; O’Neill and Felson, 2018; Wang *et al.*, 2018). The current understanding of OA development and progression is evolving from purely a mechanical disease caused by cartilage degradation towards a combined process that is mediated by the inflammatory and immunological response (Robinson *et al.*, 2016; Shu *et al.*, 2020). The synovial fluid (SF) of arthritic joints contains mediators of the inflammatory response and products of the injury to articular tissues (Simkin, 2013), while the lymphatic system plays a critical role in resolving inflammation and overall joint homeostasis (Alitalo, 2011). Despite the importance of both the lymphatic system and SF in OA disease, their relationship in OA development and progression is still poorly understood.

SF is a viscous fluid found in the cavities of synovial joints functioning as a biological lubricant and a means for nutrients and cytokines transportation (Knox, Levick and McDonald, 1988; Levick and McDonald, 1995; Blewis *et al.*, 2007; Luisa Calich, Domiciano and Fuller, 2010). Multiple inflammatory and anti-inflammatory molecules secreted from joint tissues and discovered in the SF of diseased OA patients serve as a direct indication of their role in OA development (Grissom *et al.*, 2014; Wojdasiewicz, Poniatowski and Szukiewicz, 2014). Patients with OA demonstrate increased levels of prostaglandins (PGE2), leukotrienes (LKB4), cytokines, growth factors, and nitric oxide. The inflammatory cytokines, IL-1β, IL-6, IL-15, IL-17, IL-18, and tumor necrosis factor α (TNFα), along with the anti-inflammatory cytokines, IL-4, IL-10, IL-13, can induce cartilage degradation and collagen destruction, and have thus been directly implicated within the progression of OA (Farahat *et al.*, 1993; Melchiorri *et al.*, 1998; Massicotte *et al.*, 2002; Wojdasiewicz, Poniatowski and Szukiewicz, 2014; Mora, Przkora and Cruz-Almeida, 2018). In addition, OA patients show elevated levels of the transforming growth factor β (TGFβ), fibroblast growth factors (FGFs), nerve growth factor (NGF), and vascular endothelial growth factor (VEGF) in SF, chondrocytes, subchondral bone, and serum (Hamilton *et al.*, 2016; Nagao *et al.*, 2017; Mora, Przkora and Cruz-Almeida, 2018). Notably, the increased expression of VEGF, which is known to regulate vascular permeability and angiogenesis, indicates the potential involvement of both the blood and lymphatic vasculatures in OA disease (Hamilton *et al.*, 2016; Nagao *et al.*, 2017). Although there is historical evidence that SF composition can be used for OA early detection, development, and progression (Wilkinson and Jones, 1962), to our knowledge, there is no effective use of SF constituents for therapeutic purposes.

Joint clearance and biodistribution have traditionally been assessed using radiolabeled molecule tracking and fluorescence imaging (Whitmire *et al.*, 2012; Kim *et al.*, 2015; Pradal *et al.*, 2016). Through imaging techniques, it has been demonstrated that the rate of clearance of SF constituents from the joint space significantly alters as the arthritis developed (Simkin, 2013). The rate of loss from the synovial cavity of key SF components such as, proteoglycan subunit, hyaluronic acid (HA), inflammatory cytokines, and growth factors has been shown to be relatively slow, potentially imitating conditions of clearance and drainage through lymphatics (Bard, King and Dingle, 1987; Simkin, 2013, 2015). Molecules above a certain molecular weight appear to clear at rates independent of their size, indicating a common, bulk-flow pathway consistent with lymphatic drainage (Rodnan and Maclachlan, 1960; Brown, Cooper and Bluestone, 1969). In OA, there in evidence of increased breakdown of macromolecules like HA, which could potentially lead to elevated clearance rates, turnover, and drainage into the lymphatics (Bowman *et al.*, 2018; Gupta *et al.*, 2019). Additionally, decreased viscosity of the SF in disease, and increased permeability of the synovial membrane could lead to increased transport of SF molecules into the lymphatics in the OA disease state (Wallis, Simkin and Nelp, 1987; Mwangi *et al.*, 2018; Partain *et al.*, 2020). Notably, the known effect of age on both lymphatic function and permeability and OA disease serves as an additional indication of a potential correlation between lymphatics and OA (Kushner and Somerville, 1971; Pasquali-Ronchetti *et al.*, 1992; Zolla *et al.*, 2015). Despite the indirect linkage between SF and lymphatics, the direct effect of SF exposure on LECs and lymphatic vessel function and biomechanics is not known.

There is a growing body of evidence indicating a link between OA and the lymphatic system; though these studies have mostly consisted of observed differences on lymph flow and the rate of fluid drainage between healthy and OA knee joints (Reimann *et al.*, 1989; Walsh *et al.*, 2012; Shi *et al.*, 2014). First, Wilkinson and coworkers demonstrated the presence of lymphatic vessels (LVs) in human synovial tissue while subsequent studies showed differences in both the number and size of the lymphatic vessels found in the joints (Wilkinson and Edwards, 1991; Shi *et al.*, 2014). Mice and rat models of OA have also shown decreased clearance from the synovium (Shi *et al.*, 2013; Mwangi *et al.*, 2018), alterations in lymphatic capillary density and fewer numbers of mature LVs in the late stage of disease (Shi *et al.*, 2013). Furthermore, blocking lymphangiogenesis with VEGR3 neutralizing antibodies in a mouse model reduced synovial drainage and worsened disease progression (Wang *et al.*, 2019). Given the established implication of the lymphatic system in the disease of OA, there is an increasing interest of studies trying to target lymphatics as a therapeutic modality (Han *et al.*, 2020). Although there is evidence of a correlation between the lymphatic system and OA, its specific role and function is not yet fully explained.

Despite studies suggesting that a potential effect of SF in the lymphatic system may provide further insight in OA disease, to our knowledge no study has systemically examined the response of lymphatic cells and vessels to SF treatment *in vitro* and *ex vivo*. Here, we sought to investigate the impact of SF from OA patients and healthy individuals on lymphatic endothelial cell (LEC) migration and LV contractility. We hypothesize that pump function of lymphatic vessels draining the joint could be directly compromised due to contents within the inflamed SF and thus might present a therapeutic target in OA.

## Methods

### Endothelial cell culture

*In vitro* experiments were conducted using human LECs obtained from primary cell isolation using podoplanin selection with magnetic beads following a previously established protocol (Podgrabinska *et al.*, 2002). The cells were cultured in EBM basal medium (Lonza, CC-3121) containing supplements that include 20% FBS (R&D Systems, Minneapolis, MN; S11150), 1% penicillin-streptomycin-amphotericin B (ThermoFisher Scientific, Waltham, MA; 15240062), 1% Glutamax (ThermoFisher Scientific; 35050061), 50 μM DBcAMP (Sigma Aldrich, St. Louis, MO; D0627, and 1 mg/ mL hydrocortisone acetate (Sigma Aldrich; H4126). Cells used for experiments were between passages 7 and 11. LECs were incubated at 37 °C and 5% CO_2_.

### Scratch assay

LECs were seeded in a 24-well plate. The experiment was performed with LECs at confluency (fraction of maximum cell density) of 80% to ensure sufficient contact with neighboring cells. After reaching the desired confluency, LECs were treated with OASF#3 (+20% FBS), HSF (+20% FBS), and EBM (+20% FBS) for 24 hrs. After treatment, a single scratch was made in each confluent cell monolayer using a 200 μl pipette tip and cells were washed gently in Dulbecco’s phosphate buffered salt (DPBS; ThermoFisher Scientific; MT21030CV). As negative control, LECs were treated with EBM (0% FBS) at the time of the “scratch”, respectively. Images were captured at the beginning of the experiment (*t* = 0 hrs) and at *t* = 8 and 24 hrs to monitor cell migration. We compared the images to quantify the migration rate of the cells measuring the scratch width. The presented values were normalized based on the scratch width at *t* = 0 hrs for each corresponding condition.

### Animals and isolated vessels preparation

Male Lewis rats 350-400 g, (Charles River Laboratories, Wilmington, MA) were used for all experiments. For isolation of segments of femoral LVs, the animals were euthanized via CO_2_ gas.

For the isolation of rat femoral LV (RFLV), the skin and superficial fascia layer were quickly removed to expose the area between the saphenous artery and inguinal ligament of the internal surface of the thigh (Fig. 1). Superficial caudal epigastric artery and vein were carefully teased from adjacent tissue and kept together with inguinal lymph node (Fig. 1a). They were handled gently to prevent bleeding and placed aside to provide access to the femoral lymphatics, which are located alongside the femoral vein between the femoral and superficial caudal epigastric artery junction and iliac-femoral artery (Fig. 1b). During the dissection, the area of interest was kept moist with DPBS. Suitable LVs were found along these vessels. Sections of the RFLVs ~0.5 cm long were dissected, cleaned from connective tissue, and placed in a warm (37 °C) bath of DMEM/F12 (ThermoFisher Scientific; 11039047) that was pH adjusted to 7.4 in 5% CO_2_ incubator.

**Figure 1.**
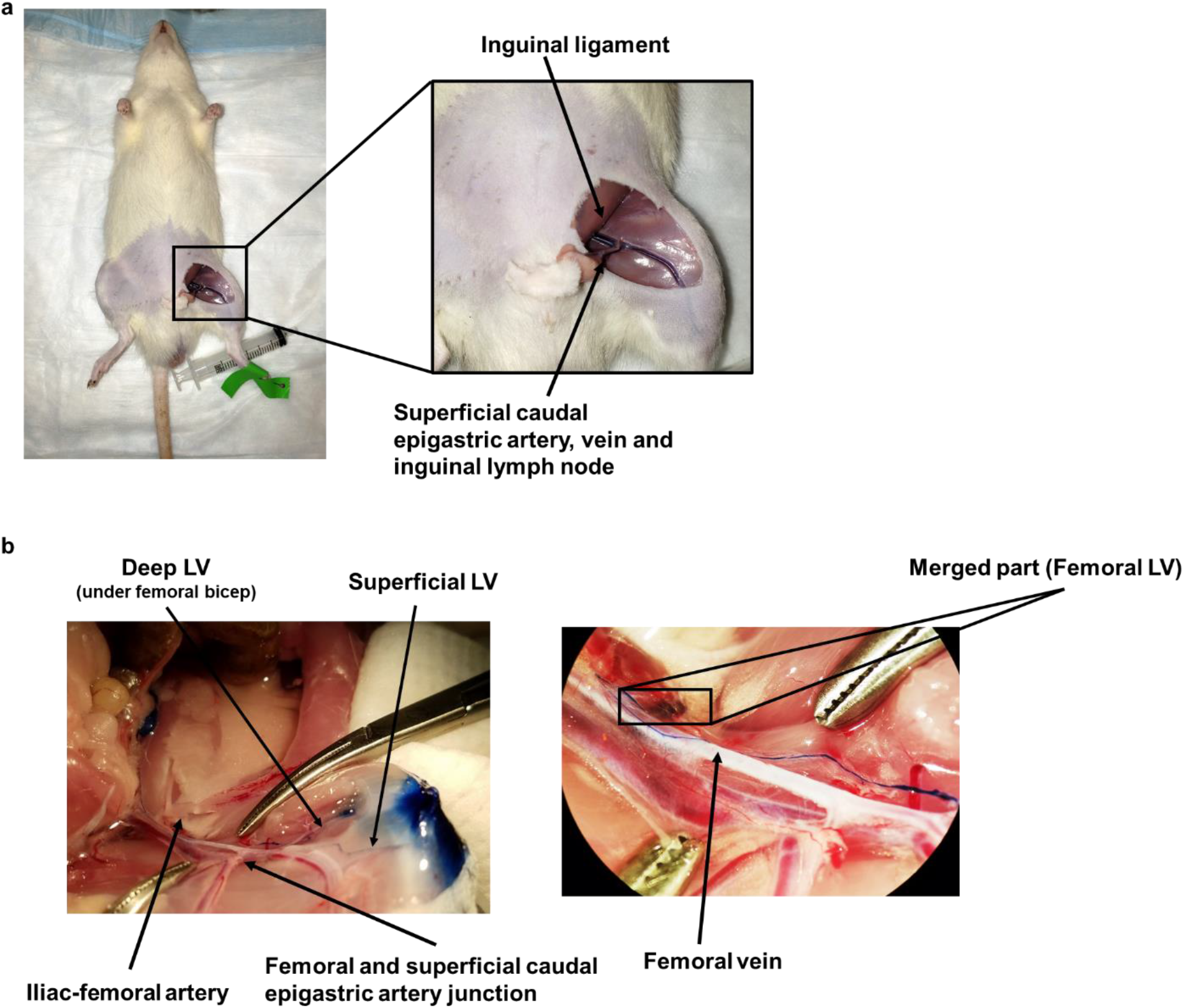
Rat femoral lymphatic vessel isolation. **(a)** Access the RFLV by sectioning the knee area. A close-up of the section is included that demonstrates the area where the RFLV resides. **(b)** Exact location of the RFLV that was isolated. Here, Evans blue (1% (w/v)) was injected intra-articularly into the left knee for visualization purposes. The immediate uptake of Evans blue by the lymphatics allows us to visualize the LV. The isolated RFLV is found proximal to where the deep LV (LV below the femoral bicep) and superficial LV merge.

In total, we examined the contractile activity of femoral LVs from 17 Lewis rats. All use of animals was in accordance with approved procedures by the Georgia Institute of Technology IACUC Review Board.

### Synovial fluid

In this study, the effect of three types of SF was investigated: healthy synovial fluid (HSF), a mixture of synovial fluid derived from various OA patients (OASF-pool), and synovial fluid derived from one specific OA patient (OASF#3). HSF was purchased from Articular Engineering (Northbrook, IL; Donor ID Number: SF-1435). OASF was collected from knee OA patients receiving arthrocentesis (joint aspiration). Patients were recruited from Emory University Sports Medicine under an Institutional Review Board (IRB) OA protocol; all patients gave informed consent. OASF was directly removed from OA patients by orthopaedic physician. SF samples were kept frozen in 1 mL aliquots at −80 °C until use.

Here, for all the presented experiments, SF was diluted in 1:10 EBM before LEC incubation (*in vitro*) or 1:10 DMEM/F12 before corresponding perfusion through the LV (*ex vivo*).

### Cytokines Assay

Cytokine content of all SF samples was quantified using a bead-based multiplex immunoassay, Luminex Cytokine/Chemokine 41 Plex Kit (EMD Millipore Corporation, Burlington, MA; HCYTMAG-60K-PX41) (Fig. 2). Median fluorescent intensity values were read using Luminex xPONENT software v 4.3 in the MAGPIX system (MAGPIX-XPON4.1-CEIVD).

**Figure 2.**
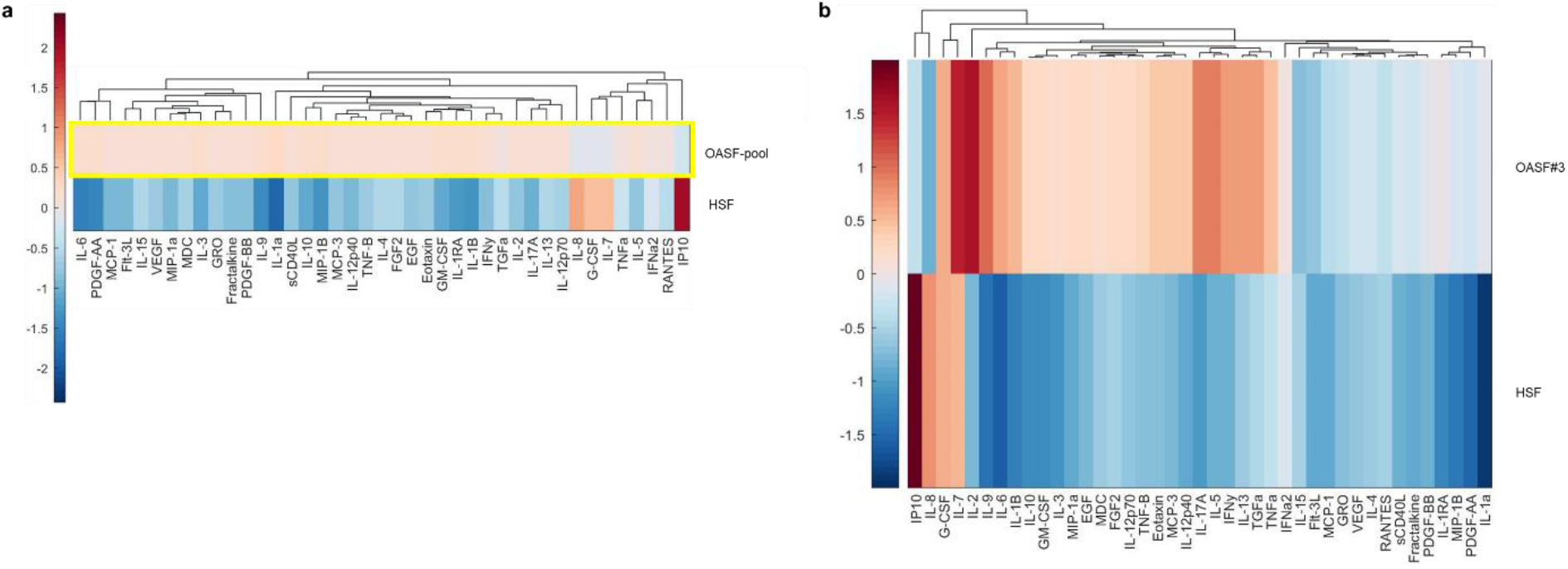
Cytokines profile for OASF and HSF. The presence of 41 markers has been assessed in **(A)** OASF-pool and HSF, and **(B)** OASF#3 and HSF using LUMINEX. The data were normalized based on the corresponding background measurement. Blue indicates a low value while red a high value.

For all SF samples, the data showed increased levels of IFN-γ inducible protein (IP-10), which has been demonstrated to be inversely associated with the severity of knee OA (Saetan *et al.*, 2011). Also, there was a common increased expression of granulocyte colony-stimulating factor (G-CSF), IL-8, and IL-7 among OASF-pool, OASF#3, and HSF, which are all shown to be implicated in pain and partial OA phenotype (Symons *et al.*, 1992; Kaneko *et al.*, 2000; Van Roon *et al.*, 2003; Ponchel *et al.*, 2005; Van Roon and Lafeber, 2008; Cornish *et al.*, 2009; Takahashi, de Andrés, *et al.*, 2015; Christensen *et al.*, 2016; Zhang *et al.*, 2016; Lee *et al.*, 2017; Sasaki *et al.*, 2017). The corresponding SF cytokines profiles were utilized along with our results to facilitate appropriate interpretation.

### Ex vivo experimental set up

An *ex vivo* lymphatic perfusion system was customized to control average transmural pressure applied to the lymphatic vessel (Fig. 3). The segment of LV without branches was cannulated on two resistance-matched glass pipettes of 150–200 μm tip diameter using the single vessel chamber (Living System Instrumentation, St Albans, VT; CH-1) (Fig. 3b). After cannulation, the chamber was connected via tubing to two glass flasks containing DMEM/F12 and two syringe pumps (Genie Touch™ Syringe Pump; Kent Scientific Corporation, Torrington, CT), and immediately placed onto the microscope stage.

**Figure 3.**
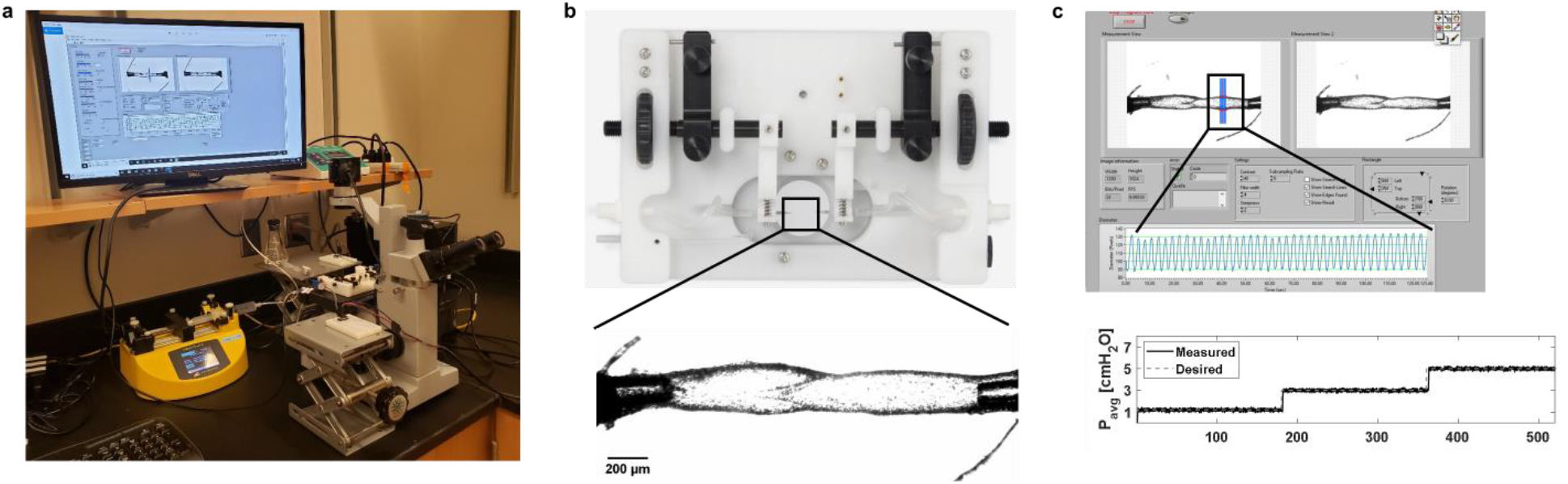
*Ex vivo* experimental set up. **(a)** Picture of the *ex vivo* lymphatic perfusion system including the computer, syringe pump, and microscope. **(d)** Picture of the single vessel chamber with a close-up of the cannulated RFLV. (5x objective; Scale bar = 200 μm.) **(c)** A custom LabView program tracks changes of the vessel diameter using data from a bright-field camera in real-time and stores this for subsequent data analysis.

The syringe pumps were used to apply the same transmural pressure to the inlet and outlet of LV. Transmural pressure was adjusted using a feedback loop with two syringe pumps and pressure sensors (1 psi SSC series; Honeywell, Charlotte, NC). During the experiment, the diameter tracing was recorded in real time via a custom LabView program, using data from a bright-field camera capturing at 30 fps, as in other studies (Gashev *et al.*, 2004) (Fig. 2c). Both diameter data and other sensor data were synchronized post-experiment using recorded time stamps. From the lymphatic diameter tracings, the following lymph pump parameters were calculated (Scallan and Davis, 2013):

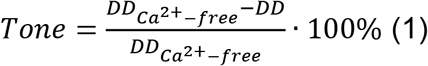

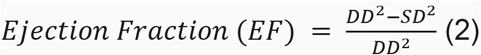

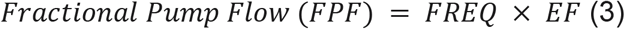

where, 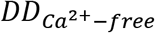 is the completely relaxed diameter as obtained with Ca^2+^ free physiological solution, *DD* the diastolic diameter, *SD* the systolic diameter, and *FREQ* the contraction frequency. The tone denotes the percentage of vessel constriction at the diastolic diameter compared to the completely relaxed diameter as obtained with Ca^2+^ free physiological solution. The ejection fraction (*EF*) indicates the part of the end-diastolic volume which was ejected during lymphatic contraction. Finally, the fractional pump flow (*FPF*) is an index of active lymph flow that produces the fractional change in lymphatic volume per minute.

For the experiments with SF treatment, the “inlet” side of the vessel was removed from the glass pipette, and SF was introduced to the corresponding upstream tubing and glass cannula. For the control experiments with DMEM/F12, the “inlet” side of the vessel was removed from the glass pipette, and DMEM/F12 was introduced to the corresponding upstream tubing and glass cannula. Then, the vessel was re-cannulated and flow with a pressure gradient of 1 cm H_2_O was generated for 5 seconds (while holding the average transmural pressure to 3cm H_2_O) to confirm successful introduction of OASF, HSF, and DMEM/F12 through the vessel. Next, the vessel containing SF or DMEM/F12 was kept at 37 °C for ~1 hr at a transmural pressure of 3 cm H_2_O and the diameter was recorded to monitor changes in contractile behavior. After the 1hr-, 2hr- and overnight treatments, average transmural pressures of 1, 3, and 5 cm H_2_O were applied to quantify the contractile response of RFLVs. At the end of each experiment, the vessel was equilibrated in Ca^2+^-free physiological solution containing (in mM): 119.0 NaCl, 4.7 KCl, 2.5 CaCl_2_, 1.2 MgSO_4_, 25.0 NaHCO_3_, 1.2 KH_2_PO_4_, 0.027 EDTA and 5.5 glucose for 20 min and subsequently exposed to an average transmural pressure of 1, 3, and 5 cm H_2_O to determine the passive diameter (for calculation of tone) at each pressure.

### Statistical analysis

For the analysis of the *in vitro* studies, a two-way ANOVA analysis was performed. For the *ex vivo* experiments, multiple comparisons were performed using a one-way ANOVA followed by a Dunnett multiple-comparison correction. For all cases, significance was defined as *p* < 0.05 (#, *, Ø) or *p* < 0.01 (##, **), *p* < 0.001 (###, ***), or *p* < 0.0001 (####, ****). For the *in vitro* studies, a # denotes a comparison with the control EBM (0% FBS) case. For the *ex vivo* experiments, a # denotes comparison with the control DMEM/F12 case, a * with the HSF case, and a Ø a comparison between the OASF-pool and the OASF#3 cases.

## Results

### Synovial fluid treatment increases the migratory response of lymphatic endothelial cells *in vitro*

We sought to investigate the effect of SF treatment on human LECs *in vitro*. The intrinsic ability of LECs to move around even in the absence of flow has been previously determined (Surya *et al.*, 2016; Michalaki *et al.*, 2020). Additionally, LECs have been shown to migrate efficiently in response to external mechanical and biomolecular cues (Kazenwadel *et al.*, 2012; Sabine *et al.*, 2012; Surya *et al.*, 2016; Michalaki *et al.*, 2020). Thus, we established a “scratch” assay to quantify cell migration (see Methods – *Scratch Assay)*. For this assay, LECs were allowed to grow to confluence and were treated either with EBM (0% FBS), OASF#3 (+20% FBS), HSF (+20% FBS), or EBM (+20% FBS). A scratch across LEC monolayers was made and LEC migration in response to designated treatments was measured following 8 and 24 hrs (Fig. 4a). A quantification based on normalized scratch width revealed that statistically significant LEC migration was achieved after treating the LECs with OASF#3 (+20% FBS), HSF (+20% FBS), and EBM (+20% FBS) compared to the EBM (0% FBS) case after 24 hrs (Fig. 4b). Notably, SF treatments led to similar phenotypes compared with EBM treatment of LECs, which corresponds to the normal cell culture medium used to maintain the LEC line. Together, although SF treatments increase the migratory response of LECs *in vitro*, they do not lead to a distinguishable phenotype from the physiologically EBM treated LECs.

**Figure 4.**
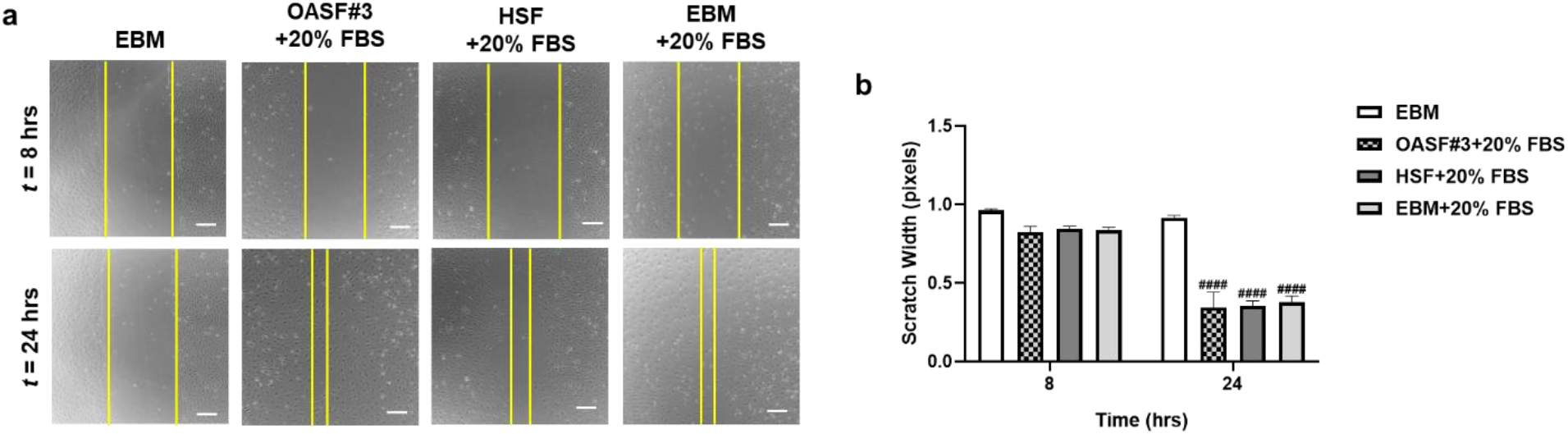
SF treatment increases LEC migration *in vitro*. HLECs treated either with EBM (0% FBS), OASF#3 (+20% FBS), HSF (+20% FBS), or EBM (+20% FBS). **(a)** The scratch in the HLEC monolayer was monitored at *t* = 8, 24 hrs. (4x objective; Scale bar = 100 μm.) **(b)** Normalized scratch width at *t* = 8 and 24 hrs. Experiments were performed in triplicate. All data represent mean values, and the error bars correspond to the standard error of the mean for each condition. Symbols on top of error bars denote comparisons using two-way ANOVA with, *p* < 0.0001 (####) vs. EBM.

### OA synovial fluid treatment transiently enhances the tonic contractions of rat femoral lymphatic vessels

Given the indistinguishable migratory effect of SF treatment groups on LECs *in vitro*, next we sought to investigate the effect of SF treatment on RFLV contractile behavior *ex vivo*. A single RFLV was isolated and cannulated on the *ex vivo* lymphatic perfusion system as described above (see Materials and Methods - *Animals and isolated vessels preparation* & *Ex vivo experimental set up*). Here, we treated the RFLVs with OASF#3 and HSF, similar to the *in vitro* studies, and also included treatment with a mixture of OASF (SF pooled from six OA patients; OASF-pool).

First, we sought to reveal the direct effect of SF treatment on RFLV diameter. To do so, we tracked the raw traces of RFLV contractions during the pressure step protocols after 1 hour and overnight treatments (Fig. 5 & Fig. S1). We found that longer incubation times lead to a substantial decrease in lymphatic contractility. Notably, OASF treatment (both OASF-pool and OASF#3) generated either an abrupt reduction after 1 hr of treatment or a complete cease of contractions after an overnight treatment, while HSF treatment displayed a gradual decrease in lymphatic contractility. For the DMEM/F12 case, RFLV seemed to remain functional as indicated by the presence of frequent diameter oscillations.

**Figure 5.**
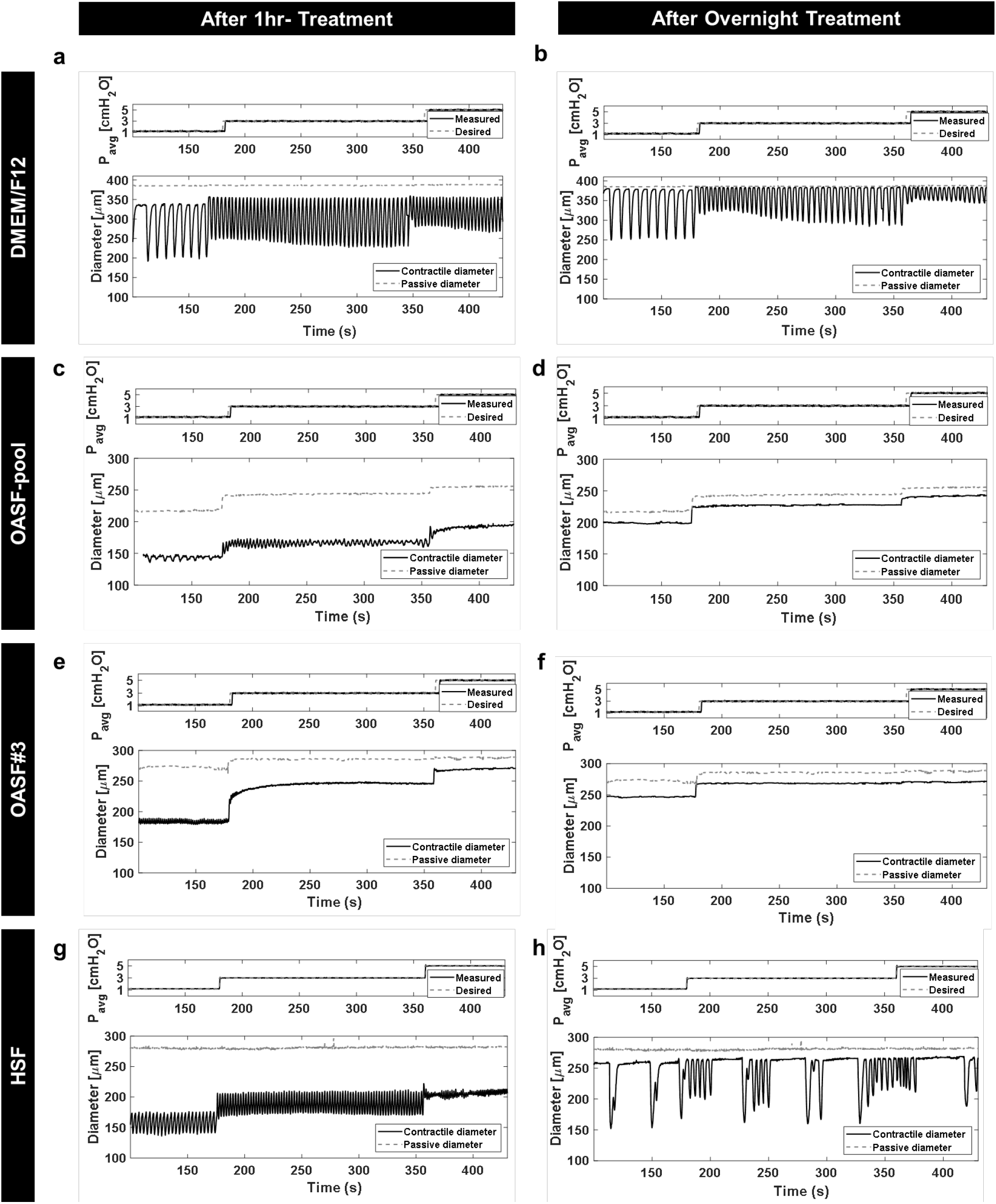
OASF treatment leads to the abrupt decrease of RFLV contractions. Diameter tracing: **(a), (c), (e), (g)** After 1hr- treatment, and **(b), (d), (f), (h)** After overnight treatment. Individual plots display trace diameters of RFLVs in: **(a, b)** DMEM/F12 (*n*=7), **(c, d)** OASF-pool (*n*=4), **(e, f)** OASF#3 (*n*=3), and **(g, h)** HSF (*n*=3). Average input and output pressures (cm H_2_O) are displayed on the top trace of each plot. They were changed simultaneously from 1 to 3 and 5 cm H_2_O (labelled). The outer diameter (μm) was measured continuously over time and plotted on the bottom trace (solid line). The outer diameter in Ca^2+^-free physiological solution (passive diameter; dashed line) was measured at the end of each experiment and plotted above the contractile diameter.

Next, we sought to examine the effect of SF treatments on the tonic contractions of the isolated vessels. To do so, RFLVs were either left untreated (DMEM/F12) or treated with OASF-pool, OASF#3, and HSF for 1 hr (Fig. 6a). After 1 hour, vessels treated with OASF-pool, exhibited significantly increased tone at all three levels of transmural pressure compared to the untreated (DMEM/F12) case, while vessels treated with OASF#3 showed increase only at 1 cm H_2_O and vessels treated with HSF showed increased tone at 1 and 3 cm H_2_O. Specifically, OASF-pool significantly elevated RFLV tonic contractions (by 25.66±5.62 at avP-1, 23.42±8.48 at avP-3, and 17.02±9.09 at avP-5; *p* = 0.0005, *p* = 0.0026, and *p* = 0.013, respectively). OASF#3 led to significant increase of RFLV tonic contractions only for the lowest transmural pressure tested (by 21.25±6.01 at avP-1; *p* = 0.0201). Finally, HSF increased the tonic contraction of RFLVs in comparison to the control DMEM/F12 case (by 20.64±8.89 at avP-1 and 15.78±7.08 at avP-3; *p* = 0.0044 and *p* = 0.0374, respectively). Together, OASF-pool treatment was the only treatment that consistently led to statistically significant increased tone for all applied transmural pressures.

**Figure 6.**
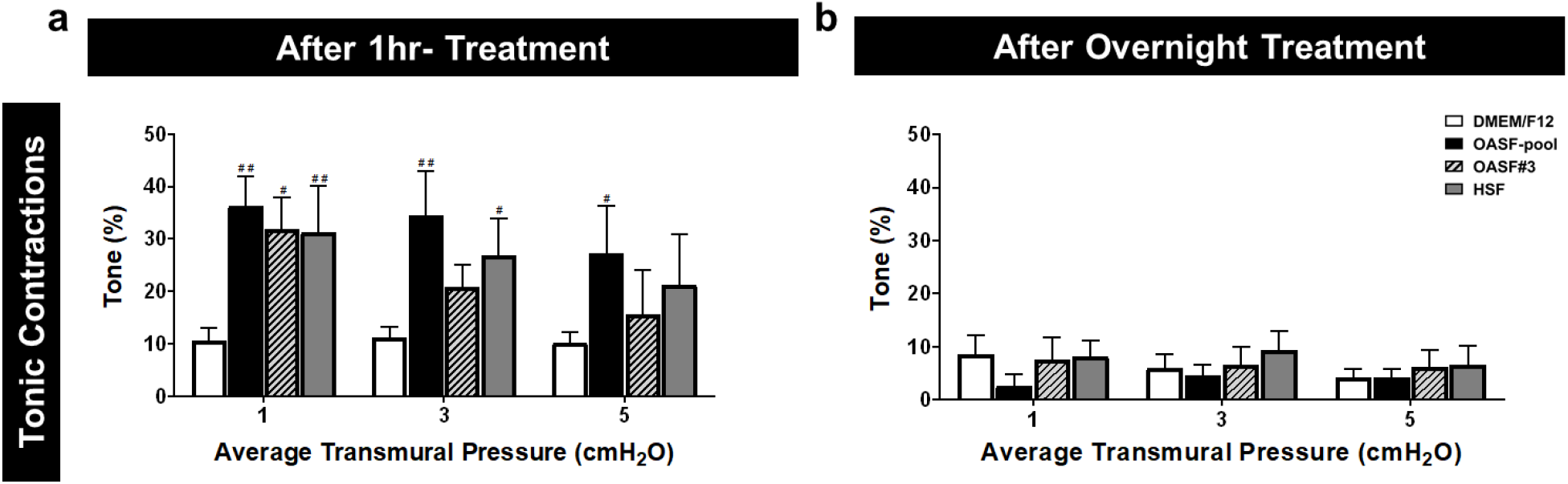
OASF treatment transiently enhances the RFLVs tonic contractions. Plots of tonic contractions. Tone (%) after treatment for **(a)** 1hr and **(b)** overnight. Each plot displays data from experiments at average transmural pressures of 1, 3, and 5 cm H_2_O for RFLVs in DMEM/F12 (*n*=7; white color), OASF-pool (*n*=4; black), OASF#3 (*n*=3; lined grey), and HSF (*n*=3; dark grey). All data represent mean values, and the error bars correspond to the standard error of the mean for each condition. Symbols on top of error bars denote comparisons using one-way ANOVA followed by a Dunnett multiple- comparison correction with, *p* < 0.05 (#) vs. control DMEM/F12.

Next, we investigated whether this enhanced tonic contraction observed during the short SF treatment (1hr- treatment) was maintained after a longer treatment (overnight treatment) (Fig. 6b). We found that tone was still present (around 5-10%) and was within the level of untreated control conditions regardless the presence of treatment with no significant differences observed. We also incubated RFLVs with SF for 2 hrs (Fig. S2a). We found that the trend of the 2hrs-treatment of the enhanced tonic contraction observed at one hour began to be diminished. Specifically, at the highest transmural pressure (avP-5) the tone of all the conditions (untreated and treated) remained low and were not statistically different from untreated vessels (Fig. S1a). Also, we found that at intermediate transmural pressure (avP-3) only OASF-pool treatment led to a significant change on tonic contractions. For low transmural pressure (avP-1), all SF treatments led to statistically significant tone index elevation compared to the DMEM/F12 case.

### OA synovial fluid treatment reduces the phasic contractions of rat femoral lymphatic vessels

Here, we set out to investigate the effect of SF treatment on phasic contractions, namely the strong periodic contractions from diastolic to systolic diameter of the LV as can be seen in Fig. 3. After treatment for 1 hour, we found that the various SF treatments yielded different behaviors on the metrics quantifying pump function, namely frequency, ejection fraction, and fractional pump flow (Fig. 7a-c). Specifically, both OASF-pool and OASF#3 decreased the frequency of lymphatic contraction at the intermediate transmural pressure (by 10.78±4.27 and 14.02±3.25 at avP-3, respectively; *p* = 0.0088 and *p* = 0.0012, respectively), while it completely stopped the phasic contractions at higher transmural pressure (at avP-5). Contrarily, HSF increased the frequency of lymphatic contractions compared to the control DMEM/F12 case for intermediate and high average transmural pressures (by 6.08±2.08 at avP-3 and 6.25±2.43 at avP-5; *p* = 0.005 and *p* = 0.0218, respectively). Regarding the ejection fraction and fractional pump flow, SF treatment led to similar results. OASF-pool decreased the ejection fraction of lymphatic contractions compared to the control DMEM/F12 case (by 0.53 ±0.01 at avP-1 and 0.43±0.02 at avP-3; *p* < 0.0001 and *p* < 0.0001, respectively). In addition, OASF#3 decreased the ejection fraction of lymphatic contractions compared to the control DMEM/F12 case (by 0.55±0.02 at avP-1 and 0.46±0.01 at avP-3; *p* < 0.0001 and *p* < 0.0001, respectively). Again, OASF-pool and OASF#3 stopped phasic contractions at avP-5. Similarly, HSF lowered the ejection fraction (by 0.32±0.09 at avP-1, 0.29±0.05 at avP-3, and 0.25±0.05 at avP-5; *p* = 0.0006, *p* = 0.0007, and *p* = 0.0016, respectively). Notably, treatment with OASF (OASF-pool and OASF#3) led to higher reduction of the ejection than the HSF treatment at low transmural pressure. Finally, all SF treatments decreased fractional pump flow compared to the DMEM/F12 case (namely, OASF-pool: by 5.84 ±0.16 at avP-1 and 7.26±0.27 at avP-3; *p* < 0.0111 and *p* < 0.0001, respectively; OASF#3: by 5.97±0.27 at avP-1 and 7.94±0.09 at avP-3; *p* = 0.0343 and *p* < 0.0001, respectively, and HSF: by 3.56 ±0.05 at avP-3 and 4.28 ±0.05 at avP-5; *p* = 0.0124 and *p* = 0.0035, respectively). Notably, OASF treatment led to a lower fractional pump flow than the HSF treatment. Together, we determined that short SF treatment has no significant effect on contractile frequency at low transmural pressure (avP-1), while at intermediate (avP-3) and high (avP-5) values OASF (OASF-pool and OASF#3) decreases while HSF increases the observed frequency. Notably, OASF#3 leads to statistically significant decrease of frequency compared to the OASF-pool case at intermediate transmural pressure (avP-3). Finally, all treatments (OASF and HSF) lead to substantial decrease in the fractional pump pressure for all applied pressures. Particularly, OASF treatment leads to a complete cease of contractions at high transmural pressure (avP-5).

**Figure 7.**
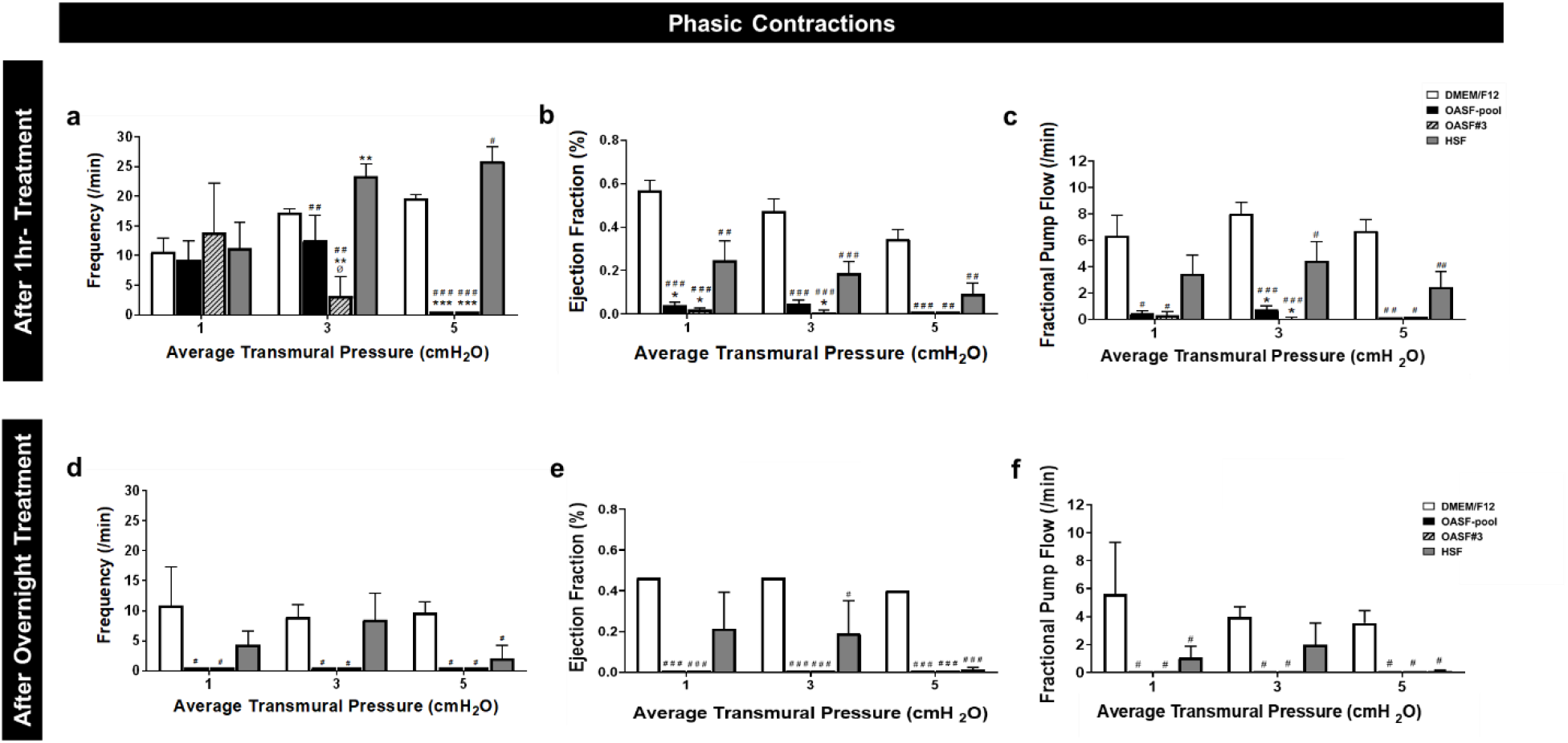
OASF treatment gradually reduces and eventually ceases the RFLVs phasic contractions. Analysis of phasic contraction metrics after treatment for **(a-c)** 1hr and **(d-f)** overnight: **(a, d)** Frequency (/min), **(b, e)** Ejection Fraction (%), and **(c, f)** Fractional Pump Flow (/min). Each plot displays data when vessels were held at average transmural pressures of 1, 3, and 5 cm H_2_O for RFLVs in DMEM/F12 (*n*=7; white color), OASF-pool (*n*=4; black), OASF#3 (*n*=3; lined grey), and HSF (*n*=3; dark grey). All data represent mean values, and the error bars correspond to the standard error of the mean for each condition. Symbols on top of error bars denote comparisons using one-way ANOVA followed by a Dunnett multiple-comparison correction with, *p* < 0.05 (#), *p* < 0.01 (##), *p* < 0.001 (###) vs. control DMEM/F12; *p* < 0.05 (*), *p* < 0.01 (**) vs. HSF; *p* < 0.05 (Ø) OASF-pool vs. OASF#3.

Finally, we sought to determine whether these phenotypes were maintained after a longer incubation (overnight treatment) (Fig. 7d-f). OASF-pool and OASF#3 completely ceased phasic contractions at all applied transmural pressures, i.e. at avP-1, avP-3, and avP-5. HSF, on the other hand, significantly decreased the frequency of contractions only at high transmural pressure (by 7.55 ±2.12 at avP-5; *p* = 0.0195). HSF significantly reduced ejection fraction (by 0.27±0.16 at avP-3 and 0.39±0.01 at avP-5; *p* = 0.0114 and *p* < 0.0001, respectively), and fractional pump flow (by 4.54± at avP-1 and 3.47±0.08 at avP-5; *p* = 0.05 and *p* = 0.0498, respectively). Together, we found that overnight treatment with HSF, while having some negative effect on lymphatic contraction, is not nearly as severe as the negative effects of OASF (OASF-pool and OASF#3) on lymphatic pump function. Experiments conducted after 2hrs-treatment followed the same trend with the overnight treatment (Fig. S2b-d).

## Discussion

Based on historic evidence, the lymphatic system has been proposed to play an important role in the development and progression of OA (Reimann *et al.*, 1989; Wilkinson and Edwards, 1991; Walsh *et al.*, 2012; Shi *et al.*, 2014). Although the involvement of lymphatics has been established for a long time, only recent studies have focused on utilizing the lymphatic system as therapeutic modality for OA (Han *et al.*, 2020). Furthermore, studies have demonstrated the role of SF as a lubricant containing multiple inflammatory and anti-inflammatory cytokines potentially crucial for OA development (Knox, Levick and McDonald, 1988; Levick and McDonald, 1995; Blewis *et al.*, 2007; Luisa Calich, Domiciano and Fuller, 2010). Furthermore, there is an established correlation between SF clearance and progression of arthritis (Simkin, 2013). Specifically, the rate of clearance of SF constituents has been demonstrated to be size-independent resembling a common pathway of lymphatic drainage (Bard, King and Dingle, 1987; Simkin, 2013, 2015). Changes in lymphatic function and permeability in the OA disease could potentially lead to increased SF transport into the lymphatics thus suggesting the use of SF as a promising future therapeutic modality (Wallis, Simkin and Nelp, 1987; Mwangi *et al.*, 2018; Partain *et al.*, 2020). Despite the importance of both the lymphatic system and SF in OA disease, their specific role has yet to be examined. To our knowledge, this work constitutes the first *ex vivo* investigation of how RFLVs respond to SF treatment in the case of OA.

Here, we found that SF from both OA and healthy joints increases the migratory response of LECs *in vitro* (Fig. 4). OASF treatment of isolated LVs *ex vivo* transiently enhances the tonic contractions, while gradually reduces and eventually ceases the RFLVs phasic contractions. (Fig. 5–7). Specifically, this intrinsic phasic contraction of the lymphatic is completely abolished after 24 hours of treatment, while the muscle of the vessel still maintains some function as observed by the presence of vessel tone even after 24 hours of treatment. It is worth noting that while treatment with healthy SF had an effect on contractions compared to culturing the vessels in just DMEM/F12, this effect was not nearly as severe when compared to OASF from both the pooled SF and the SF exclusively from patient 3. We chose to dilute the SF such that 10% of the volume of the media perfused through the lumen of the vessel contained SF. This was an approximation as it is not clear what the exact dilution of SF contents is within the pre-nodal lymph immediately draining the vessel. Furthermore the relative concentrations of proteins and molecules between the two compartments is likely a function of size as has been shown in other tissue beds (Levick and Mcdonald, 1995; Miller *et al.*, 2011). In addition, it is worth noting that the frequency of lymphatic contractions in these idealized isolated vessel preparations is often higher than what is observed *in vivo*, suggesting that there are factors from the *in vivo* context that are not recapitulated in the isolated vessel preparation.

While the exact molecular mechanisms responsible for this reduction in lymphatic contraction remain unknown, data from the cytokine array provides insight into molecules that are elevated and warrant further investigation (Fig. 2). First, IFN-γ inducible protein (IP-10) levels in SF has been demonstrated to be inversely associated with the severity of knee OA (Saetan *et al.*, 2011) demonstrating the validity of our data (high IP-10 levels *only* in HSF; Fig. 2). Through close inspection of the cytokines profile, we found there is a common increased expression of granulocyte colony-stimulating factor (G-CSF), IL-8, and IL-7 among HSF, OASF-pool, and OASF#3. Studies have suggested the use of G-CSF as a safe therapeutic strategy for arthritis and generally in inflammatory conditions where pain is important (Cornish *et al.*, 2009; Christensen *et al.*, 2016; Lee *et al.*, 2017; Sasaki *et al.*, 2017). Here, the increased levels observed in all types of SF used, implies the presence of extreme inflammatory conditions of all donors. IL-8 has been demonstrated to reach high expression levels in OA patients (Symons *et al.*, 1992; Kaneko *et al.*, 2000; Takahashi, de Andres, *et al.*, 2015). IL-7 has shown to play a crucial role in cartilage destruction during rheumatic diseases and has been correlated with the initiation and progression of OA (Van Roon *et al.*, 2003; Ponchel *et al.*, 2005; Van Roon and Lafeber, 2008; Zhang *et al.*, 2016). The high cytokine expression levels of IL-8 and IL-7 in the SF of all donors suggest the presence of a partial OA phenotype. This analysis could justify the gradual diminishment of both tonic and phasic contractions observed with the increase of SF treatment time. The *ex vivo* observations seem to oppose the *in vitro* findings, where LEC mobility seems to increase in the presence of SF while LV contractility gradually ceases. This provides strong evidence on the importance of establishing representative *ex vivo* models for the investigation of OA.

Our study possesses several limitations. One such limitation is the limited sample size of HSF and OASF-pool. Due to the sparsity of SF from healthy individuals and the limited availability of SF from OA patients, the experimental outcome is highly patient-dependent. Thus, the interpretation/ extrapolation of our observations should be conducted with caution. Another similar restriction is the limited amount of SF, which challenged the execution of both the *in vitro* and *ex vivo* experiments. Knowing the limited stock of SF accessible both in lab (OASF) and commercially (HSF), we minimized the number while simultaneously reassuring the quality of our experiments. Another limitation is the use of human-derived SF to treat human LECs (*in vitro* studies) and rat-derived LVs (*ex vivo* studies). The use of different species could justify the disparity in the observed phenotypes between the *in vitro* and *ex vivo* experiments.

Nevertheless, our work is the first attempt to utilize SF directly derived from OA patients and healthy individuals to investigate its potential effect on lymphatic contractility. The striking effect of SF, both OASF (OASF-pool and OASF#3) and HSF, on the tonic and phasic lymphatic contractions serves as a further indication of the importance of the lymphatic system in the concept of OA (Wilkinson and Edwards, 1991; Yoshida *et al.*, 1997; Bouta *et al.*, 2015; Simkin, 2015; Doan *et al.*, 2019). A potential use of SF as a means of therapeutic for OA should be further investigated.

