## Supplemental Figures for "Effect of human synovial fluid from osteoarthritis patients and healthy individuals on lymphatic contractility"

### Supplemental Material

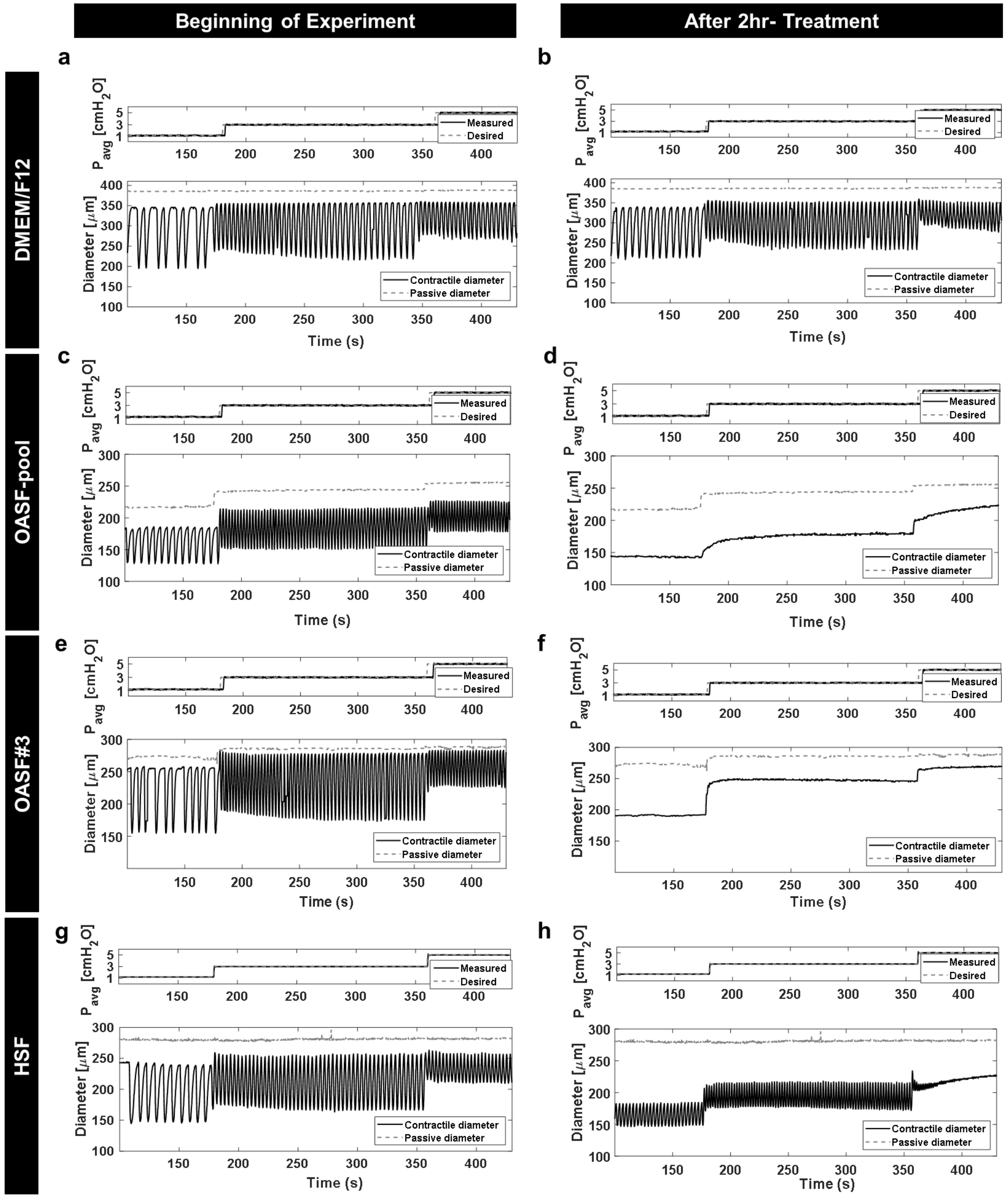

**Figure S1. Raw traces of RFLV contractions during the pressure step protocol in DMEM/F12, OASF-pool, OASF#3, and HSF.** Diameter tracing: **(a), (c), (e), (g)** At the beginning of the experiment and **(b), (d), (f), (h)** After 2hr- treatment. Individual plots display trace diameters of RFLVs in: **(a, b)** DMEM/F12 (*n*=7), **(c, d)** OASF-pool (*n*=4), **(e, f)** OASF#3 (*n*=3), and **(g, h)** HSF (*n*=3). Average input and output pressures (cm $H_2O$) are displayed on the top trace of each plot. They were changed simultaneously from 1 to 3 and 5 cm $H_2O$ (labelled). The outer diameter (μm) was measured continuously over time and plotted on the bottom trace (solid line). The outer diameter in $Ca^{2+}$-free physiological solution (passive diameter; dashed line) was measured at the end of each experiment and plotted above the contractile diameter.

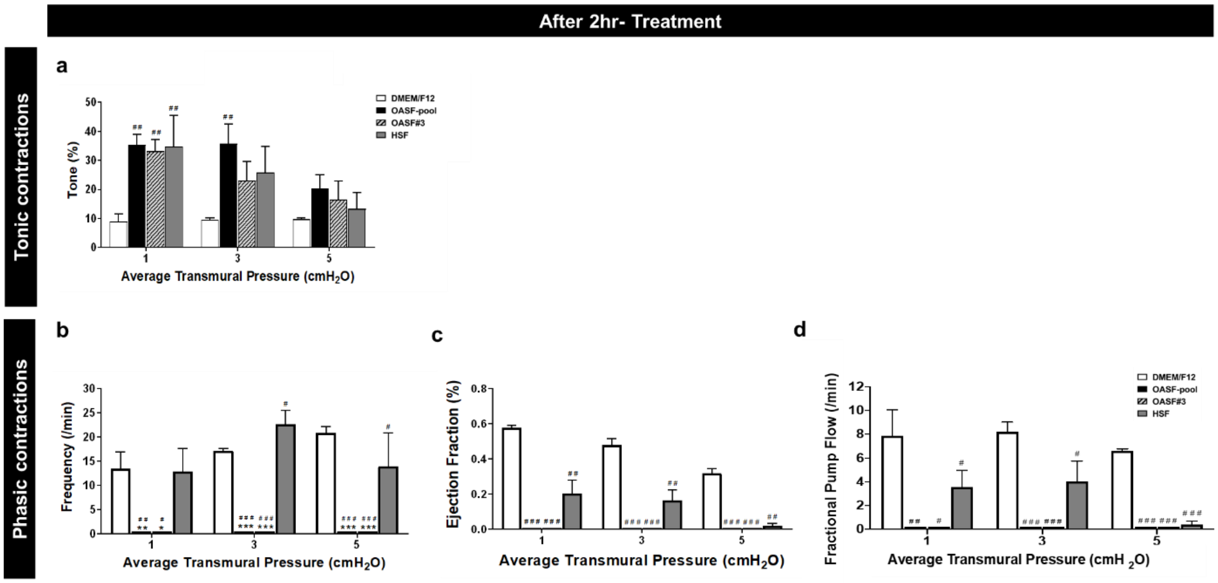


**Figure S2. Effect of DMEM/F12, OASF-pool, OASF#3, and HSF on RFLV contractility.** Plots of tonic and phasic contractions after 2hr-treatment: **(a)** Tone (%), **(b)** Frequency (/min), **(c)** Ejection Fraction (%), and **(d)** Fractional Pump Flow (/min). Each plot displays average transmural pressure of 1, 3, and 5 cm $H_2O$ for RFLVs in DMEM/F12 (*n*=7; white color), OASF-pool (*n*=4; black), OASF#3 (*n*=3; lined grey), and HSF (*n*=3; dark grey). All data represent mean values, and the error bars correspond to the standard error of the mean for each condition. Symbols on top of error bars denote comparisons using one-way ANOVA followed by a Dunnett multiple-comparison correction with, $p <$ 0.05 (#), $p < 0.01$ (##), $p < 0.001$ (###) vs. control DMEM/F12; $p < 0.05$ (*), $p < 0.01$ (**), $p < 0.001$ (***) vs. HSF.
